# MC-Bayes: A Python-based wrapper for MotionCor3 processing of EER files compatible with Bayesian polishing

**DOI:** 10.64898/2026.08.06.743412

**Authors:** Raymond N. Burton Smith, Kazuyoshi Murata

## Abstract

Here, we present MC-Bayes, a Python-based script for processing cryo-electron microscopy EER movies on one or more GPUs using MotionCor3 in a user-friendly manner. Further, it generates the .star files necessary for RELION to perform Bayesian polishing (a.k.a.: reference-based motion correction) with EER movies. Until now, Bayesian polishing of EER data was only possible if the CPU-based “RELIONCor” implementation of MotionCor2 was used, which is sub-optimal on GPU-heavy cryo-EM processing systems. This wrapper was created for those facilities and/or users who may have (many) powerful GPUs, but for whatever reason have few CPU cores or less system RAM. Leveraging MotionCor3, MC-Bayes allows motion correction of EER data 2 or more times faster (depending on system) than the RELION CPU implementation, except in circumstances where dozens or hundreds of CPU cores with high quantities of system RAM can be utilised.

## Introduction

With the development of Electron Event Representation (EER) (Guo et al., 2020) and its implementation as output from Thermo Fisher Scientific Falcon 4/4i cameras, both temporal and spatial flexibility post-acquisition were placed in the hands of cryo-EM users. This wrapper puts back in the hands of those using the EER format with RELION the speed of GPU accelerated motion correction: MotionCor2 (Zheng et al., 2017) and 3 (Zheng, 2024) do not output the files required for RELION (Fernandez-Leiro & Scheres, 2017; Kimanius et al., 2016; Scheres, 2012; Zivanov et al., 2018; Zivanov et al., 2019, 2020) to carry out Bayesian polishing when EER format input is given, and RELION’s implementation (Zivanov et al., 2018) is CPU-based and a poor choice for those using GPU-heavy processing systems. Here, we show that our wrapper out-performs the CPU-based RELIONCor implementation on a cryo-EM workstation which has a good general balance between CPU and GPU.

MC-Bayes was designed to have as few dependencies as possible. It has two: `tifffile`, so that EER fraction counting can be performed and the internal fraction grouping can be calculated, and `tqdm`, for live feedback on progress of motion correction. The RELION 5 and 5.1 Python environments both contain the required dependencies. MC-Bayes is the product of our testing the utility of local LLM models (Qwen3-coder-next (Cao et al., 2026) and Qwen-3.6-35B-A3B) to integrate and improve a set of private bash scripts which perform the same functions but are independent and must be run in order (and are also rather user-unfriendly).

Using this wrapper on GPU heavy systems, one of the early bottlenecks of EER image processing is removed, showing RELION remains a powerful tool for fast image processing of biological cryo-EM single particle analysis (SPA).

## Results

By default, the wrapper (https://github.com/rbs-sci/MC-Bayes) asks a series of questions about the basic nature of the data, automatically detects a gain reference, assigns valid nVIDIA GPUs and begins processing. If a . `star` file is specified (e.g., from a RELION Import job) it can automatically read essential electro-optical parameters unless overridden (see Appendix 1 for a description of command line arguments). Figure 1 shows the questions posed and information output when a . `star` file is provided.

**Figure 1:**
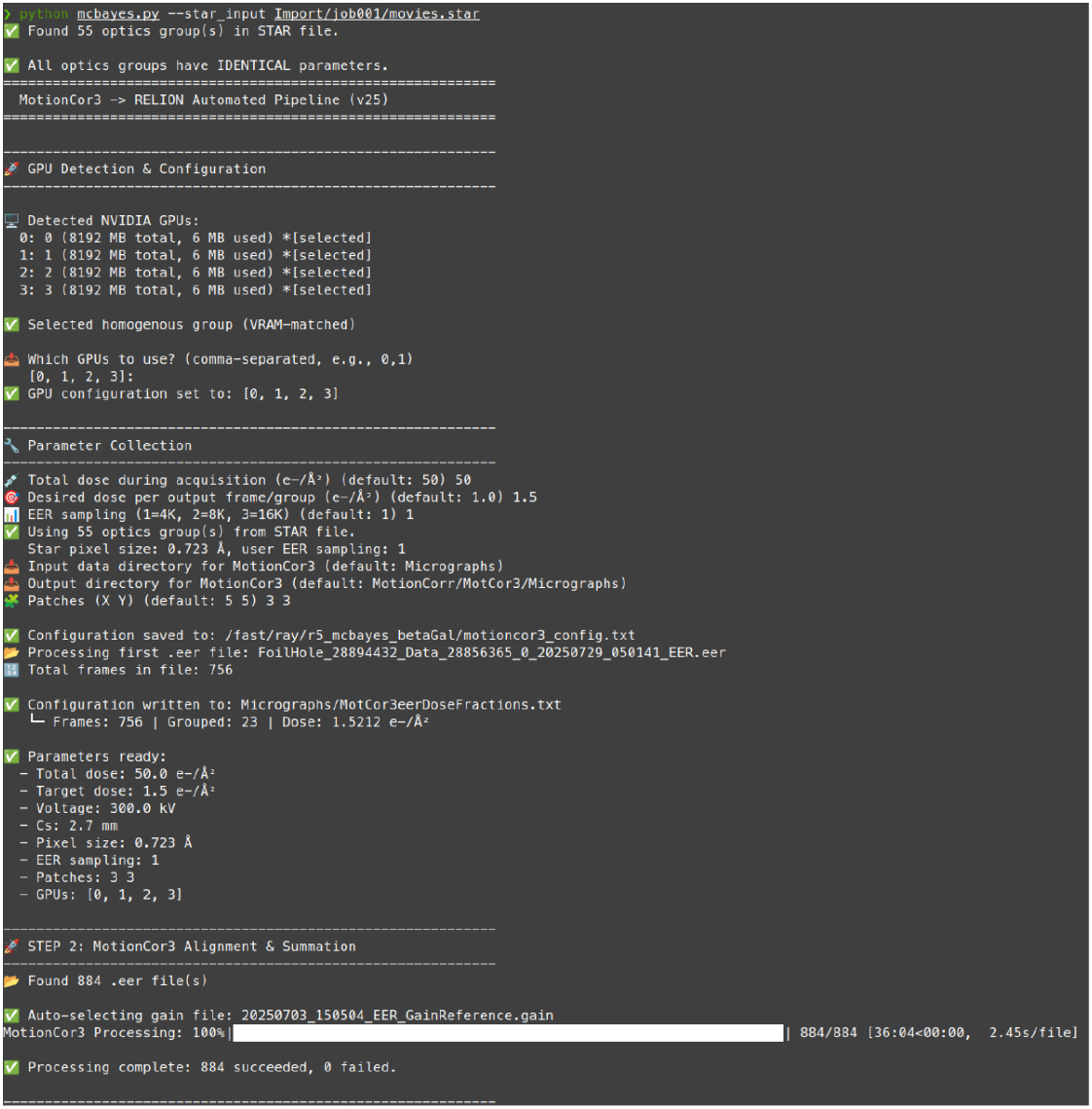
Screenshot of the MC-Bayes parameter acquisition interactive session.

The wrapper outputs both `micrographs_corrected.star` (for passing to CTFFIND (Rohou & Grigorieff, 2015)) and `micrographs_ctf.star` (as MotionCor3 calculates CTF parameters on the GPU) for immediate curation and particle picking. In our testing, CTFFIND results in superior CTF estimates – more micrographs have good CTF fits – regardless of whether the power spectrum output or micrograph output is used for CTF estimation when compared to the estimates from MotionCor3 and should be used.

To reduce filesystem performance having an impact on the comparison, movies were copied to local PCI-E SSD storage. To avoid file RAM caching influencing read speed, the system was rebooted between each run, whether MotionCor3 wrapper or RELIONCor. The test system was an AMD Threadripper Pro 5975WX (32 core/64 thread) with 512GB of RAM (DDR4 at 3200 MT/s) equipped with 6x nVIDIA RTX A4000 (Ampere generation) cards. This was chosen as while is has many GPUs, it also has a surfeit of RAM and a moderate core count, meaning RELIONCor can be run in a way to provide best performance: many MPI (message passing interface) processes. For this testing, 16 MPI processes were used for CPU-based motion correction and was performed with and without Hyper-Threading (HT), as many shared-access clusters disable this for security reasons (Table 1). HT reduced time required for CPU motion correction ∼33% when doubling number of threads per MPI process, although this requires further testing. Table 1 contains timings on a few systems tested: on an older system with 128GB of system RAM, multi-GPU MC-Bayes provides a clear speed advantage over RELIONCor (2.6× faster) and adding further MPI processes results in running out of memory. On another system, 22,700 EER micrograph movies collected with the same parameters as the β-galactosidase dataset described below were motion corrected across 10 GPUs in approximately 10 hours (data not shown) when streamed from network storage.

**Table 1:** Some statistics for time taken for motion correction with RELIONCor and the MotionCor3 MC-Bayes wrapper on two systems of varying specifications.

| Dataset | Apoferri<br>itin | β-<br>galactosi<br>dase | β-<br>galactosi<br>dase | β-<br>galactosi<br>dase | β-<br>galactosi<br>dase | β-<br>galactosi<br>dase | β-<br>galactosi<br>dase |
| --- | --- | --- | --- | --- | --- | --- | --- |
| System | 1 | 1 | 1 | 2 | 2 | 2 | 2 |
| CPU cores | 32/64 | 32/64 | 32/64 | 8/16 | 8/16 | 8/16 | 8/16 |
| RAM<br>(GB) | 512 | 512 | 512 | 128 | 128 | 128 | 128 |
| GPU<br>(Type) | A4000<br>(Amp) | A4000<br>(Amp) | A4000<br>(Amp) | RTX<br>2080 | RTX<br>2080 | RTX<br>2080 | RTX<br>2080 |
| GPU (No.) | 6 | 6 | 6 | 4 | 4 | 4 | 4 |
| Pixel size | 0.57 | 0.723 | 0.723 | 0.723 | 0.723 | 0.723 | 0.723 |
| Total dose | 50 | 50 | 50 | 50 | 50 | 50 | 50 |
| Dose/frame | 0.9404 | 0.7937 | 1.5873 | 0.97 | 0.97 | 0.97 | 0.97 |
| No. mics | 1160 | 884 | 884 | 884 | 884 | 884 | 884 |
| Time (MC-Bayes) (mm:ss) | 23:24 | 21:46 | 22:08 | 0:36:04 | - | - | - |
| Time (per mic) (s) | 1.21 | 1.48 | 1.5 | 2.45 | - | - | - |
| RELIONC or MPI Processes | 16M4T | 16M4T | 16M4T | - | 4M4T | 8M2T | 16M1T |
| Time (RELION Cor) (no HT) (h:mm:ss) | 1:06:16 | 1:02:02 | - | - | - | - | - |
| Time (RELION Cor) (HT) (h:mm:ss) | 0:47:20 | 0:47:35 | 0:47:52 | - | 1:46:20 | 1:36:00 | CRASH |

Two datasets were tested for direct comparison, subsets of internal data and long-standing benchmark samples: apoferritin and β-galactosidase. We were focussed on whether MC-Bayes can match or better the results from RELIONCor: in terms of speed, it is comparable to CryoSPARC (Punjani et al., 2017) Patch Motion with equivalent parameters on the same system. Very occasionally, MotionCor3 will get “stuck” on a micrograph, taking significantly longer (>1 minute) than expected to process it. This seems to happen more on older GPUs (Turing/RTX 2000 and older) and we have not yet been able to clearly isolate a cause.

The `eer_trajectory_handler` script provided with RELION which allows resampling/regrouping of EER fractions also functions correctly with wrapper output - at time of writing, this is an advantage over CryoSPARC EER handling, where resampling is not yet available.

### Apoferritin

Since 2014 and the availability of direct detectors, resolving the structure of apoferritin by cryo-EM became a reality (Russo & Passmore, 2014). Because of its robustness and high symmetry, it routinely gives high-resolution results, it has become the new de facto standard for cryo-EM (Danev et al., 2026; Hamaguchi et al., 2019; Nakane et al., 2020; Yip et al., 2020). Except in exceptional circumstances, it is the only biological complex to have achieved true atomic resolution with cryo-EM which many laboratories have now demonstrated. 1,160 micrograph movies, collected at 0.57 Å/pixel using EPU and beam-image-shift acquisition were used. All exposures were from from a single grid square. They were processed with RELION 5.1 using either RELIONCor (rc) (Figure 2) or MotionCor3 using the MC-Bayes wrapper (mcb) (Figure 3). Final results between the two are close to indistinguishable, but motion correction was faster with MC-Bayes (23 minutes versus 47 minutes on system 1) (Table 1). RELIONCor showed a slight advantage in reported b-factor (Table 2).

**Table 2:** Image processing information for the four primary runs.

| | rc apoferritin | mcb apoferritin | rc β-galactosidase | mcb β-galactosidase |
| --- | --- | --- | --- | --- |
| Micrographs | 1,160 | 1,160 | 884 | 884 |
| Passed CTF estimation (<6 Å) | 1154 | 1154 | 867 | 868 |
| Picked particles | 192,528 | 215,020 | 128,746 | 130,820 |
| Selected particles | 179,335 | 194,620 | 107,986 | 106,265 |
| Final resolution (GS-FSC)(GS) | 1.6 | 1.6 | 1.77 | 1.74 |
| Final b-factor | -35.8 | -37.8 | -22.5 | -22.2 |
| Map-to-model FSC | 1.61 | 1.61 | 1.8 | 1.78 |

**Figure 2:**
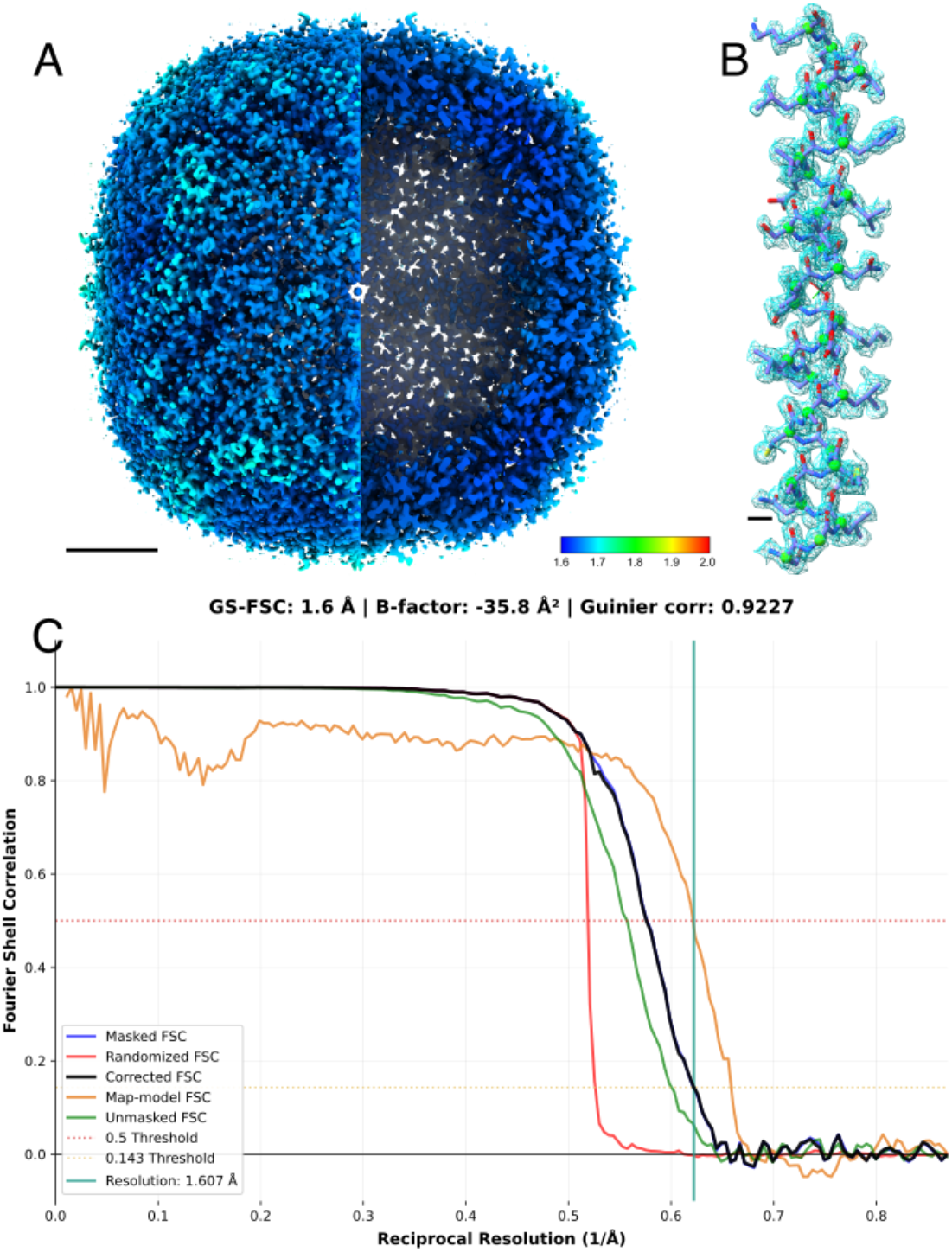
Apoferritin processed with RELIONCor. (A) Final map, coloured by local resolution, with one semi-hemisphere cut away for internal visualisation. Scale bar 2 nm. (b) A focused view of residues 95-124, with PDBID:9WAL fitted. Scale bar 2 Å. (C) Gold-standard FSC reports 1.6 Å (black line), with masked FSC shown in blue, unmasked FSC shown in green and phase randomised FSC shown in red. Map-to-model FSC calculated with a rigid-body-fit 9WAL using Servalcat.

**Figure 3:**
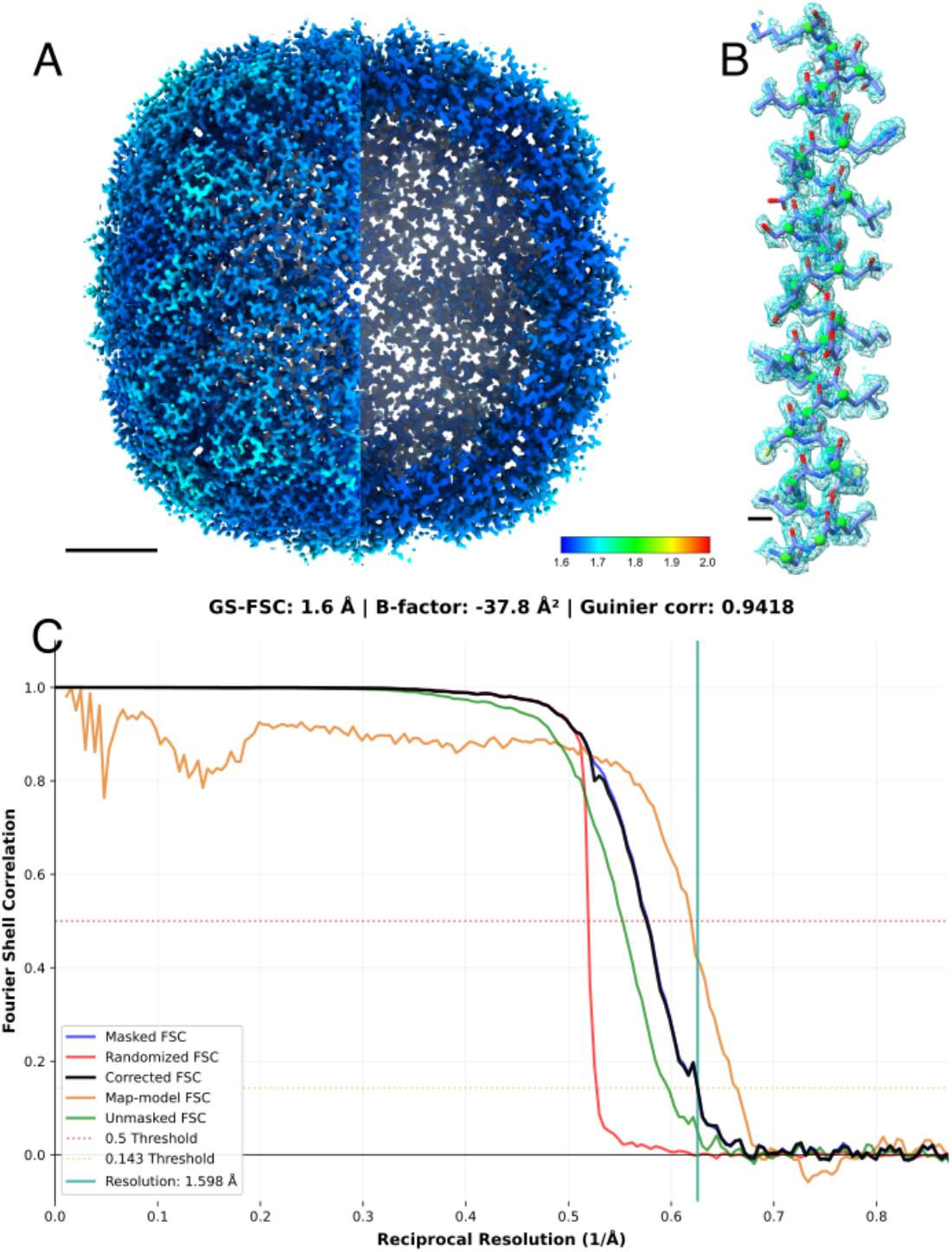
Apoferritin processed with MotionCor3 via MC-Bayes. (A) Final map, coloured by local resolution, with one semi-hemisphere cut away for internal visualisation. Scale bar 2 nm. (b) A focused view of residues 95-124 with PDBID:9WAL fitted. Scale bar 2 Å. (C) Gold-standard FSC reports 1.6 Å (black line), with masked FSC shown in blue, unmasked FSC shown in green and phase randomised FSC shown in red. Map-to-model FSC calculated with a rigid-body-fit 9WAL using Servalcat.

### β-galactosidase

A classic cryo-EM standard sample (Bartesaghi et al., 2015; Fujita et al., 2023; Kayama et al., 2021). 884 micrograph movies, collected at 0.723 Å/pixel using EPU and beam-image-shift acquisition were used. All exposures were from a single grid square. They were processed with RELION 5.1 using either RELIONCor (Figure 4) or MotionCor3 using the MC-Bayes wrapper (Figure 5). With MC-Bayes, motion correction was completed in approximately 22 minutes versus 47 minutes. Prior to CTF refinement, the mcb data was 0.2 Å higher resolution than the rc data. After CTF refinement, the reported resolutions differed by a single Fourier shell (2.16 vs. 2.17 Å) and ∼0.25 b-factor after post-processing. After Bayesian polishing, resolutions were identical and difference in b-factor remained unchanged. After further CTF refinement, they differed by three Fourier shells in favour of mcb. Final b-factors were so close for the difference to be functionally meaningless (Figs. 4C, 5C) (Table 2).

**Figure 4:**
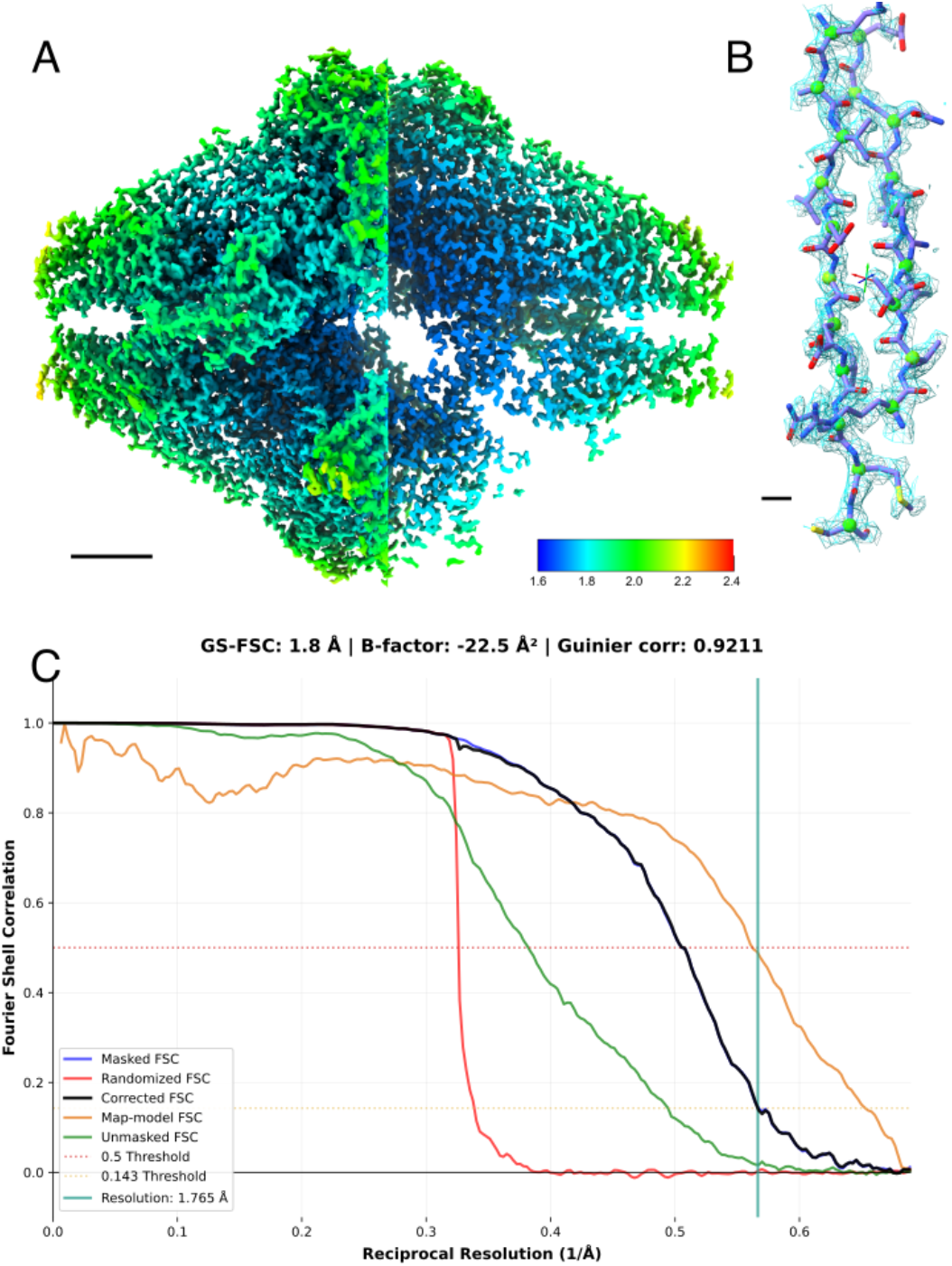
β-galactosidase processed with RELIONCor. (A) Final map, coloured by local resolution, with one semi-hemisphere cut away for internal visualisation. Scale bar 2 nm. (b) A focused view of two β-strands (residues 238-248 and 289-297). Scale bar 2 Å. (C) Gold-standard FSC reports 1.77 Å (black line), with masked FSC shown in blue, unmasked FSC shown in green and phase randomised FSC shown in red. Map-to-model FSC (yellow) calculated with a rigid-body-fit 43LS using Servalcat .

**Figure 5:**
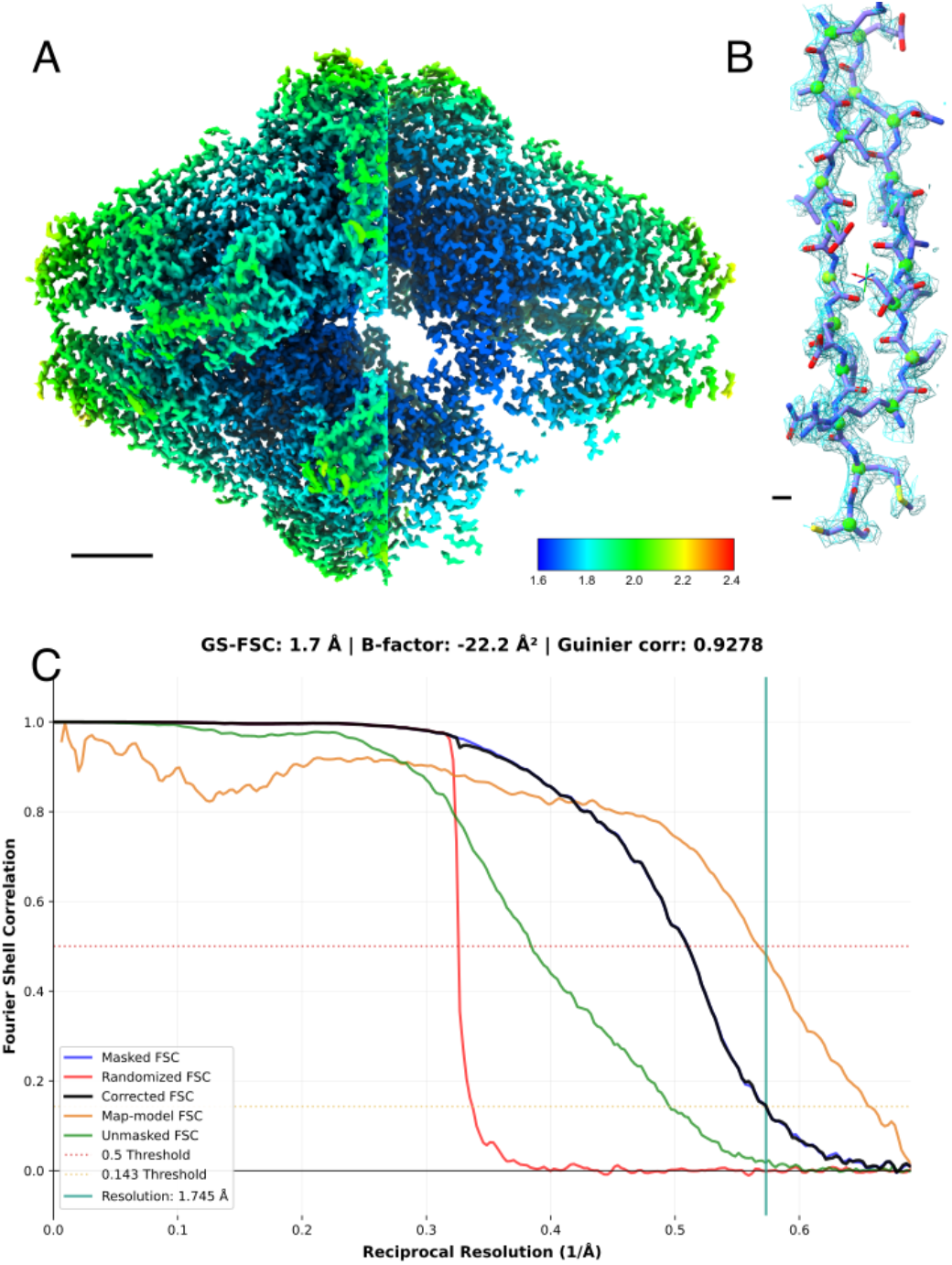
β-galactosidase processed with MotionCor3 via MC-Bayes. (A) Final map, coloured by local resolution, with one semi-hemisphere cut away for internal visualisation. Scale bar 2 nm. (b) A focused view of two β-strands (residues 238-248 and 289-297). Scale bar 2 Å. (C) Gold-standard FSC reports 1.74 Å (black line), with masked FSC shown in blue, unmasked FSC shown in green and phase randomised FSC shown in red. Map-to-model FSC (yellow) calculated with a rigid-body-fit 43LS using Servalcat .

1Discussion

For two standard samples, whether using RELIONCor or MC-Bayes as a MotionCor3 wrapper yielded near indistinguishable final reconstructions. Simultaneously, by leveraging MotionCor3 for GPU-accelerated motion correction rather than the CPU-accelerated RELIONCor, faster EER motion correction is possible for GPU-heavy cryo-EM systems whose users wish to utilise RELION rather than CryoSPARC.

For example, in our institute, we have other systems which are heavily biased toward GPU compute and perform sub-optimally with RELIONCor as a result: e.g. 18 core (36 thread) systems with 256GB RAM and quadruple RTX A5000 GPUs and 16 core (32 thread) systems with 384GB RAM and quadruple RTX A5000 GPUs. On these systems, RELIONCor creates a severe bottleneck in processing. Recommendations of converting EER to TIFF destroys the value of collecting micrographs in EER, and while it may be a solution for some, the loss of flexibility in the event that a dataset achieves better than anticipated results is a non-starter for us and many others. MC-Bayes removes this conundrum.

Of course, for TIFF or MRC files, this wrapper is unnecessary, as while MotionCor2 is no longer officially supported by its developers, MotionCor3 itself can be used as a drop-in replacement by RELION and outputs correct information for further (Bayesian polishing) processing.

We hope others who wish to continue to take advantage of the temporal and spatial flexibility offered by the EER file format, while wanting to use RELION, will find this wrapper of use.

## Methods

### Software setup

The MotionCor3 source code (v.1.2.4) was downloaded from GitHub (Zheng, 2024) and compiled with `–fPIC –fPIE` for compatibility with modern Linux kernel ASLR (address space layout randomisation) security. MotionCor3 was compiled against CUDA 12.6 using gcc 11.4 and 13.3. A fork with the appropriate modifications is available at: https://github.com/rbs-sci/MotionCor3_fPIC

### Data collection

Exposures were acquired using a Thermo Fisher Scientific Titan Krios G4 equipped with a Selectris-X energy filter and Falcon 4i direct detector at the ExCELLS, Okazaki, Japan. EPU was used for data collection, using beam image shift (“faster” mode) with default image shift range. All data were saved as EER format movies. QuantiFoil R1.2/1.3 copper grids were used. β-galactosidase was acquired at 165,000×, equivalent to 0.723 Å/pixel, with a total dose of 50 e^-^/Å^2^. Apoferritin was acquired at 215,000×, equivalent to 0.57 Å/pixel, with a total dose of 50 e^-^/Å^2^.

### Image processing

#### Apoferritin

1,160 movies were imported into RELION (Fernandez-Leiro & Scheres, 2017; Kimanius et al., 2016; Scheres, 2012; Zivanov et al., 2018; Zivanov et al., 2019, 2020) 5.1 with optics groups based on beam image shift. Motion Correction was carried out either with MotionCor3 using the MC-Bayes wrapper, or RELIONCor, using the same parameters. CTF estimation was carried out using CTFFIND (4.1.14) (Rohou & Grigorieff, 2015). Table 2 summarises statistics. Topaz picking with a diameter of 120 Å was carried out on a 100 micrograph subset, and particles extracted with a figure of merit (FoM) minimum of 0. After 2D classification using the expectation-maximisation (EM) algorithm, classes which displayed clear features were selected and the particles passed to Topaz to train a picking model. This model was then used on the whole dataset. Particles were extracted into 384 pixel boxes downsampled to 96 pixels, and 2D classified using the VDAM algorithm. Clear classes were selected and passed to initial model generation. The Blush regulariser (Kimanius et al., 2023) was not used. A 3D refinement was carried out, then particles re-extracted into 384 pixel boxes downsampled to 320 pixels. 3D refinement and CTF refinement (Zivanov et al., 2020) were carried out, then Bayesian polishing (Zivanov et al., 2019) employed, resampling to 384 pixels. After Bayesian polishing, another 3D refinement was carried out, followed by further CTF refinement. A final 3D refinement was followed by post-processing and local resolution estimation. With Ewald sphere correction (Russo & Henderson, 2018), the final resolution for both was 1.6 Å (1 d.p).

#### β-galactosidase

884 movies were imported into RELION (Fernandez-Leiro & Scheres, 2017; Kimanius et al., 2016; Scheres, 2012; Zivanov et al., 2018; Zivanov et al., 2019, 2020) 5.1 with optics groups based on beam image shift. Motion Correction was carried out either with MotionCor3 using the MC-Bayes wrapper, or RELIONCor, using the same parameters. CTF estimation was carried out using CTFFIND (4.1.14) (Rohou & Grigorieff, 2015). Table 2 summarises statistics. Topaz picking with a diameter of 180 Å was carried out on a 100 micrograph subset, and particles extracted with a figure of merit (FoM) minimum of -1. After 2D classification using the expectation-maximisation (EM) algorithm, classes which displayed clear features were selected and the particles passed to Topaz to train a picking model. This model was then used on the whole dataset. Particles were extracted into 512 pixel boxes downsampled to 128 pixels, and 2D classified using the VDAM algorithm. Clear classes were selected and passed to initial model generation. The Blush regulariser (Kimanius et al., 2023) was used for all 3D refinements. A 3D refinement was carried out, then particles re-extracted into 512 pixel boxes downsampled to 384 pixels. 3D refinement and CTF refinement (Zivanov et al., 2020) were carried out, then Bayesian polishing (Zivanov et al., 2019) employed, resampling to 512 pixels. After Bayesian polishing, another 3D refinement was carried out, followed by further CTF refinement. A final 3D refinement was followed by post-processing and local resolution estimation. Ewald sphere correction (Russo & Henderson, 2018) made no difference to the final reported resolution. Final resolution was 1.8 Å (1 d.p) for RELIONCor and 1.7 Å (1 d.p) for MotionCor3 with MC-Bayes.

### Model building

ChimeraX (Goddard et al., 2018) was used to align models and maps as appropriate. PDBID:9WAL was used for apoferritin. 43LS was used for β-galactosidase. Servalcat (Yamashita et al., 2021) was used to calculate map-to-model FSCs.

### Visualisation

Figures were made with ChimeraX (Goddard et al., 2018) (focused views with the ISOLDE (Croll, 2018) plugin). FSC curves were plotted with PrettyFSC (https://github.com/rbs-sci/prettyfsc). Figures were assembled with Inkscape.

## Data availability

Maps, half-maps and FSC curves will be made available upon reasonable request, or upon requirement by a journal.

The MC-Bayes wrapper is available at: https://github.com/rbs-sci/MC-Bayes. The PrettyFSC script is available at: https://github.com/rbs-sci/prettyfsc.

The current released version of MotionCor3 does not support operating systems which require Position Independent Code and Position Independent Executable builds, and two of the pre-compiled libraries are compiled without this support. A modified MotionCor3 repository can be found at https://github.com/rbs-sci/MotionCor3_fPIC. For convenience, download the repository and run the zz_compile.sh script to build libraries and MotionCor3. This assumes CUDA 12.6 is installed in `/usr/local/`. Adjust as appropriate.

## Appendix 1: Command line Options

MC-Bayes is run with the following syntax:

`$ python mc-bayes.py [--arguments] [Directory]`

or to use defaults and read essential parameters from RELION, use:

`$ python mc-bayes.py --star_input Import/job001/movies.star`

Help

`-h, --help:` explains what options are available in the command line

### Input

`[no input]:` directory must be defined by the user in the interactive section. Defaults to Micrographs

`[directory]:` script takes directory defined on the command line as location of raw movies. Also searches for appropriate gain reference (.`gain` or . `mrc`) in the same directory

`--star_input:` define a . `star` format file (respects optics groups, takes pixel size, spherical aberration, acceleration voltage and amplitude contrast from .star file

`--gain_file:` if you want to specify a non-standard gain file location, or a specific gain file out of several options (in latter case, script will identify multiple gain files and ask which you want to use)

`--gain_ref:` alias for `--gain_file`, because sometimes muscle memory takes over

### Output

`--output_star:` if you want your combined . `star` file named something non-standard. Defaults to `micrographs_ctf.star`

`--force:` overwrite already existing output (use with care!)

### Configuration

`--config:` load a configuration from a file (use if reproducing a run, or after config file generation with `--write-config`

`--write-config:` output a configuration file, which can be manually adjusted and then fed back in with --config

`--no-config:` do not write out a configuration file record

`--ask:` force manual parameter input even if configuration file exists

### GPU handling

`--gpus:` define which GPUs to use (follows nvidia-smi numbering)

`--heterogeneous:` use all (nVIDIA) GPUs, regardless of type/VRAM. Default: false

`--min_free_mb:` define what the minimum memory should be for the auto-selection routine to choose a GPU. Default: 4096

### Parameters

`--kv:` acceleration voltage (supports 100, 120, 200, 300). Default: 300

`--cs:` spherical aberration (in mm). Default: 2.7

`--angpix:` Pixel size in Angstrom/pixel. Default: 0.73

`--total_dose:` what total dose was applied to the sample (in e^-^/Å^2^). Default: 50

`--target_dose:` what dose-per-frame you would like. Default: 1

`--eer_sampling:` whether you want EER sampling 1 (4K output), 2 (8K output) or 3 (16K). RELION does not support 16K Bayesian polishing! 16K is for completeness (it’s still possible to break both physical and super resolution Nyquist with 16K sampling). Default: 1

`--patches:` for patch motion correction. Set, “1 1” to carry out full-frame motion correction only. Needs two numbers, space separated. Numbers do not have to be identical, but for a square sensor it makes the most sense. Default: 5 5

### Advanced

`--advanced:` expose advanced parameters which are usually OK to leave alone

`--quiet:` don’t bother me with verbose output for post-Motion Correction .star file generation

`--bft:` b-factor blurring to try to prevent alignment of high frequency noise. Default: 150

`--iter:` number of iterations to perform of alignment. Not usually necessary to adjust. Default: 10

`--tol:` convergence tolerance. Not usually necessary to change. Default: 0.1

`--ft_bin:` Fourier cropping of output. Used to be common to Fourier crop super resolution data back to physical pixel size but is now less common. Non-integer Fourier cropping CAN be used but make sure the resulting micrograph size is an EVEN NUMBER! Default: 1

`--throw:` discard this many frames from the start of the (fraction grouped) stack. Default: 0

`--trunc:` discard this many frames from the end of the (fraction grouped) stack. Default: 0

`--ampcont:` Amplitude contrast. RELION default: 0.1. MotionCor3 default: 0.07. Lots of discussion about this historically which we won’t get into here. Default: 0.1

`--fmref:` reference frame for alignment. Default: first frame

`--rotgain:` rotate gain reference clockwise in 90 degree steps. Default: 0

`--flipgain:` flip gain horizontally (along y axis, 2) or vertically (along x axis, 1) or not (0). Vertical flip required for EPU . `gain` files. Default: 1

`--invgain:` Invert gain (divide rather than multiply gain reference). Required for some early EER data. Usually recommend generating an .mrc gain using cisTEM `sum_all_eer_files` or RELION’s `relion_estimate_gain` rather than doing this

`--infmmotion:` enable or disable the in-frame motion model of MotionCor3. Meant to compensate for motion trapped within single frames. Have not tested extensively

`--tilt:` define tilt axes, if required. Default: “0 0”

`--defect_file:` provide a four-column text file listing defective pixels. Not usually necessary

`--dark_ref:` a dark reference file (.mrc format). Not usually necessary

`--init_dose:` if there was a pre-exposure dose applied to the area the micrograph(s) were taken from. Not usually necessary unless data is exotic

`--group:` per-frame grouping for motion correction. Two space separated numbers, where the first is whole frame grouping and second is patch frame grouping. Not usually necessary to change, unless using excessively fine dose fractionation where SNR is too low for accurate alignment. Default: “1 4”

## Appendix 2: Installation

The RELION 5.x Python environments run the MC-Bayes wrapper without issues. Alternatively:

~~~
$ conda create –n mcbayes python=3.14
$ conda activate mcbayes
$ pip install tifffile tqdm
~~~

Done!

To run, MotionCor3 executable (recommend latest version, 1.2.4 at time of writing) should be in the RELION project directory. It’s only small, so should pose no issues. Micrograph movies and gain reference should (ideally) be in a directory named “Micrographs”. Symlink(s) are fine.

